# Control of Munc13-1 Activity by Autoinhibitory Interactions Involving the Variable N-terminal Region

**DOI:** 10.1101/2024.01.24.577102

**Authors:** Junjie Xu, Victoria Esser, Katarzyna Gołębiowska-Mendroch, Agnieszka A. Bolembach, Josep Rizo

## Abstract

Regulation of neurotransmitter release during presynaptic plasticity underlies varied forms of information processing in the brain. Munc13s play essential roles in release via their conserved C-terminal region, which contains a MUN domain involved SNARE complex assembly, and control multiple presynaptic plasticity processes. Munc13s also have a variable N-terminal region, which in Munc13-1 includes a calmodulin binding (CaMb) domain involved in short-term plasticity and a C_2_A domain that forms an inhibitory homodimer. The C_2_A domain is activated by forming a heterodimer with the zinc-finger domain of αRIMs, providing a link to αRIM-dependent short- and long-term plasticity. However, it is unknown how the functions of the N- and C-terminal regions are integrated, in part because of the difficulty of purifying Munc13-1 fragments containing both regions. We describe for the first time the purification of a Munc13-1 fragment spanning its entire sequence except for a flexible region between the C_2_A and CaMb domains. We show that this fragment is much less active than the Munc13-1 C-terminal region in liposome fusion assays and that its activity is strongly enhanced by the RIM2α zinc-finger domain together with calmodulin. NMR experiments show that the C_2_A and CaMb domains bind to the MUN domain and that these interactions are relieved by the RIM2α ZF domain and calmodulin, respectively. These results suggest a model whereby Munc13-1 activity in promoting SNARE complex assembly and neurotransmitter release are inhibited by interactions of the C_2_A and CaMb domains with the MUN domain that are relieved by αRIMs and calmodulin.

## Introduction

Neurons communicate through neurotransmitters that are released from presynaptic terminals through Ca^2+^-evoked synaptic vesicle exocytosis and bind to receptors on the postsynaptic plasma membrane. Neurotransmitter release involves a series of steps that include tethering of synaptic vesicles to specialized areas of the presynaptic plasma membrane called active zones, priming to a release-ready state(s) and very fast (< 1 ms) fusion of the vesicle and plasma membranes upon Ca^2+^ influx.^1^ Each of these steps can be regulated by a variety of molecular mechanisms during short- and long-term presynaptic plasticity processes that result in distinct release probabilities, shaping the properties of neural networks and underlying diverse forms of information processing in the brain.^2,3^ Thus, presynaptic terminals do not act merely as passive signal transducers but in fact constitute minimal computational units in the brain.

The sophisticated protein machinery that controls neurotransmitter release and its regulation has been extensively characterized, which has led to reconstitution of basic features of synaptic vesicle fusion with purified components^4–6^ and definition of their functions (reviewed in refs.^7,8^). The neuronal SNAP receptors (SNAREs) syntaxin-1, synaptobrevin and SNAP-25 form a tight four helix bundle called the SNARE complex^9–11^ that brings the synaptic vesicle and plasma membrane together and mediates membrane fusion^12,13^. The SNARE complex is disassembled by N-ethylmaleimide sensitive factor (NSF) and soluble NSF adaptor proteins (SNAPs) to recycle the SNAREs,^9,14^ whereas Munc18-1 and Munc13s organize SNARE complex assembly via an NSF-SNAP-resistant mechanism^4,15,16^ that is initiated with Munc18-1 bound to a self-inhibited ‘closed’ conformation of syntaxin-1.^17,18^ Subsequently, Munc18-1 also binds to synaptobrevin, forming a template for SNARE complex formation^19–21^ while Munc13 bridges the vesicle and plasma membranes^22,23^, and opens syntaxin-1.^24–26^. The resulting partially assembled SNARE complex binds to synaptotagmin-1,^27^ the Ca^2+^ sensor that triggers fast release,^28^ and to complexin,^29^ forming a spring-loaded macromolecular assembly^30,31^ that is ready for fast membrane fusion upon Ca^2+^ influx through an as yet unclear mechanism.^7^

In addition to being essential for neurotransmitter release because of their role in SNARE assembly, Munc13s act as master regulators of release in a variety of presynaptic plasticity processes though their multidomain architecture.^3,32–34^ These large (ca. 200 kDa) proteins from the active zone are characterized by a large C-terminal region that is sufficient to rescue the total abrogation of neurotransmitter release observed in Munc13-1/2 double knockout mice^22^ and includes: a C_1_ domain that is responsible for diacylglycerol (DAG) and phorbol ester-dependent augmentation of neurotransmitter release;^35,36^ a C_2_B domain that binds Ca^2+^ and PIP_2_, and mediates Ca^2+^-dependent short-term presynaptic plasticity;^37^ a MUN domain^38^ that mediates syntaxin-1 opening;^24,25^ and a C_2_C domain that binds membranes^23^ (see domain diagram for Munc13-1, the most abundant mammalian Munc13 isoform, in Figure 1A). The crystal structure of a Munc13-1 fragment spanning the C_1_, C_2_B and MUN domains (referred to as CCM, Figure 1A)^39^ revealed how the C_1_ and C_2_B domains, which are expected to bind to DAG and PIP_2_ at the plasma membrane, emerge at one end of the highly elongated MUN domain, while the other end should be attached to the C_2_C domain (which was not in the structure but can be modeled with AlphaFold^40^; Figure 1B). These findings suggested that Munc13-1 bridges synaptic vesicles to the plasma membrane through respective interactions with the C_2_C domain and the C_1_/C_2_B region,^22,39^ which was supported by diverse evidence and was shown to be critical for neurotransmitter release^23^. Moreover, the Munc13-1 C-terminal region is believed to mediate such bridging in two distinct orientations that allow distinct extents of SNARE complex assembly^39^, and the more slanted orientation, which is more active, is most likely favored by Ca^2+^ binding to the C_2_B domain and DAG binding to the C_1_ domain^41^. This orientation switch was proposed to underlie the Ca^2+^- and DAG-dependent enhancement of neurotransmitter release mediated by the Munc13-1 C_2_B and C_1_ domains, respectively, which was supported by mutagenesis and electrophysiological data.^41^ Importantly, a model invoking two primed states with different release probabilities and distinct Munc13-1 orientations can explain a large amount of available presynaptic plasticity data.^42,43^

**Figure 1.**
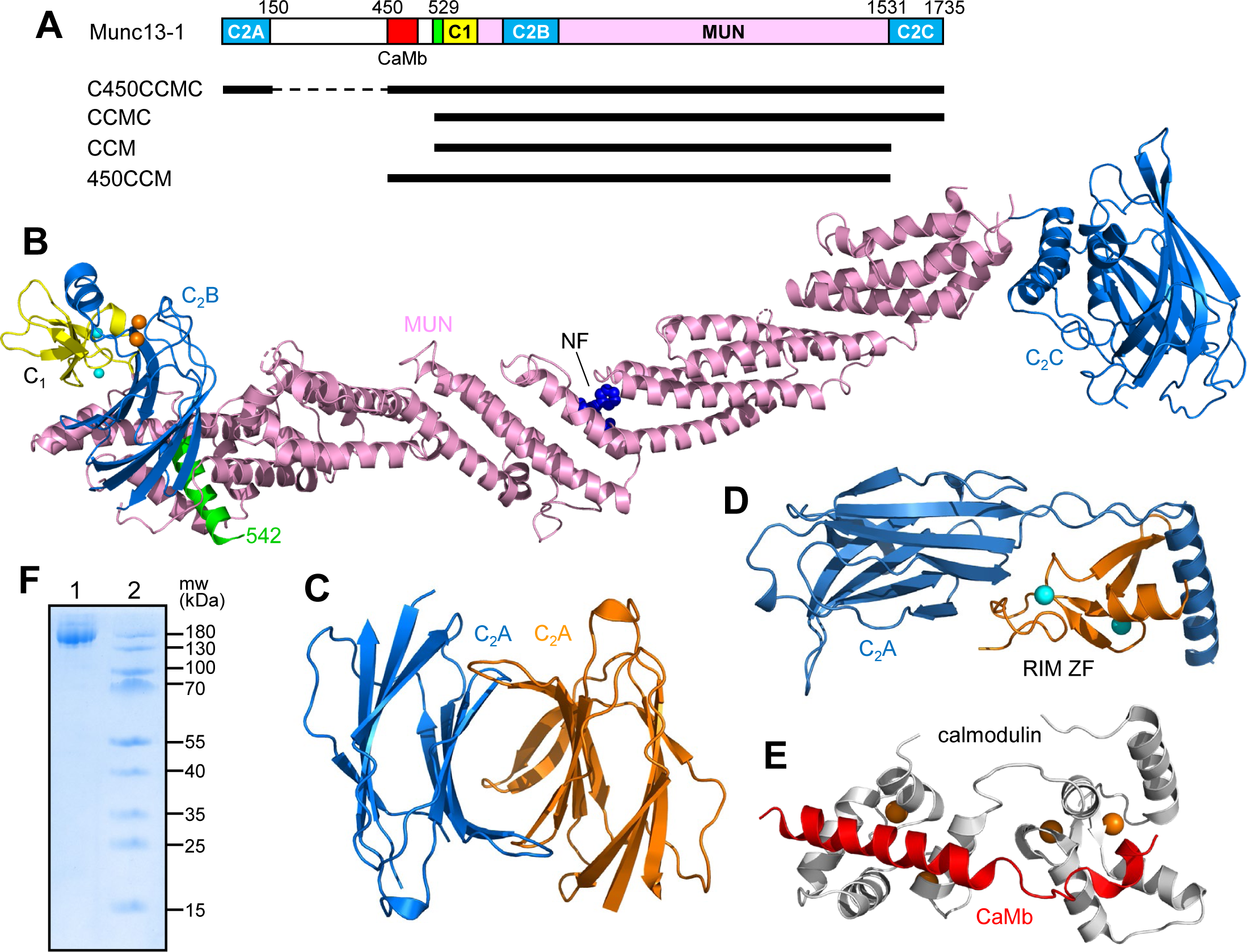
Structures of Munc13-1 domains and analysis of C450CCMC purity. (A) Domain diagram of Munc13-1 and summary of large Munc13-1 fragments used in this study. Residue numbers corresponding to approximate domain boundaries are indicated above the diagram. The black bars below the diagram illustrate the regions of Munc13-1 included in each fragment, with the abbreviation used for designation of each fragment on the left. (B) Ribbon diagram of a model of the conserved C-terminal region of Munc13-1, with the N-terminal helix in green, the C_1_ domain in yellow, the MUN domain in pink and the C_2_B and C_2_C domains in blue. Ca^2+^ and Zn^2+^ ions are shown as orange and cyan spheres, respectively. Residues Asn1128 and Phe1131, which are important for the activity of the MUN domain in opening syntaxin-1,^25^ are shown as dark blue spheres. The model was built with the crystal structure of the CCM fragment^39^, completing the Ca^2+^-binding region of the C_2_B domain with the crystal structure of this domain bound to Ca^2+^ ^37^ (PDB IDs 5UE8 and 3KWU, respectively), and with a model of the C_2_C domain produced by AlphaFold^40^ (https://alphafold.ebi.ac.uk/). The orientation of the C_2_C domain with respect to the MUN domain is based on the different orientations observed in the AlphaFold model of Munc13-1 and cryo-EM crystal structures of the Munc13-1 C-terminal^62^, but this orientation is likely flexible given its variability in the different structures. The position of the most N-terminal residue observed in the crystal structure of the CCM fragment (residue 542) is indicated to show how the sequences N-terminal to these residues are expected to be oriented toward the MUN domain. (C, D) Crystal structures of the Munc13-1 C_2_A domain homodimer (C) and the heterodimer of the Munc13-1 C_2_A domain (blue) with the RIM2α ZF domain (orange)^44^ (D) (PDB IDs 2CJT and 2CJS, respectively). (E) NMR structure of calmodulin (gray) bound to the Munc13-1 CaMb sequence (red)^52^ (PDB ID 2KDU). (F) SDS-PAGE analysis with Comassie blue staining of purified C450CCMC (lane 1). Molecular weight markers are in lane 2 and their molecular weights are indicated on the right.

Munc13s also contain a variable N-terminal region that in Munc13-1 includes (Figure 1A): a C_2_A domain that forms a homodimer^44^ and a heterodimer with the N-terminal zinc finger (ZF) domain of αRIMs,^45,46^ which are large Rab3 effectors that organize the active zone and mediate some forms of short-term plasticity as well as mossy fiber long-term potentiation;^47,48^ and a calmodulin binding domain (CaMb) that mediates short term Ca^2+^-dependent plasticity^49^ and is separated from the C_2_A domain by a low-complexity sequence. Homodimerization of the Munc13-1 C_2_A domain inhibits neurotransmitter release and formation of the heterodimer with αRIMs releases this inhibition,^50^ helping to localize Munc13-1 at the active zone to mediate vesicle priming.^45,51^ These observations indicate that Munc13-1-αRIM binding provides a means to couple diverse forms of Rab3- and RIM-dependent presynaptic plasticity to the core apparatus that mediates synaptic vesicle priming and fusion. Intriguingly, deletion of the Munc13-1 C_2_A domain strongly impairs synaptic vesicle priming and neurotransmitter release, whereas deletion of the entire N-terminal region has much milder effects, and deletion of residues 150-520, which span the low-complexity sequence and the CaMb domain, does not substantially alter vesicle priming or Ca^2+^-evoked release.^51^ These findings suggest that the Munc13-1 C_2_A domain also has a functional interplay with the CaMb domain that may connect calmodulin-dependent plasticity to Rab3- and RIM-dependent regulation of release. Although the available three dimensional structures of the Munc13-1 C_2_A domain homodimer and its heterodimer with the RIM ZF domain^44^ (Figure 1C,D), and of the complex of calmodulin with the Munc13-1 CaMb domain^52^ (Figure 1E) have revealed the nature of these binary interactions, it is still unknown how these interactions and those involving the Munc13-1 C-terminal region are integrated to coordinate the multiple functions of Munc13-1 in neurotransmitter release and presynaptic plasticity. Furthermore, models produced by AlphaFold^40^ (https://alphafold.ebi.ac.uk/) do not yield reliable information on potential intramolecular interactions of the Munc13-1 C_2_A and CaMb domains with the C-terminal region. Hence, it is unclear whether the phenotypes observed with the various deletions arise from alteration of intramolecular interactions or of binding to other factors.

Recognizing that addressing these questions requires studies of purified full-length Munc13-1 or fragments including the C_2_A domain and/or the CaMb domain in addition to the conserved C-terminal region, we initiated attempts to prepare such proteins over a decade ago, but this research was hindered by the high tendency of these large proteins to degradation. Here we describe the culmination of these efforts, which led to the expression and purification of a fragment that spans the entire Munc13-1 sequence except for the low complexity sequence (residues 151-449) and hence includes the C_2_A domain, the CaMb domain and the C-terminal region (referred to as C450CCMC). Using fusion assays with reconstituted proteoliposomes, we show that C450CCMC is much less active in promoting liposome fusion than a fragment corresponding to the C-terminal region spanning the C_1_, C_2_B, MUN and C_2_C domains (referred to as CCMC), but the activity of C450CCMC is strongly stimulated by the Rim2α ZF domain together with calmodulin. NMR and gel filtration assays show that both the C_2_A domain and the CaMb domain bind to the MUN domain and that these interactions are released by the RIM2α ZF domain and calmodulin, respectively. These results suggest that intramolecular interactions between the variable N-terminal region and the conserved C-terminal region control Munc13-1 activity and may underlie an intricate coupling between distinct forms of presynaptic plasticity dependent of Munc13-1.

## Results

### Expression and purification of Munc13-1 C450CCMC

The major problems that we had to overcome to obtain purified fragments spanning most or all of the Munc13-1 sequence after expression in bacteria were the low expression yields and the tendency of the proteins to be degraded by bacterial proteases remaining even after affinity chromatography, ion exchange chromatography and gel filtration. The strategy that we used to obtain Munc13-1 C450CCMC fragment involved the use of a double affinity tag, with a maltose-binding protein (MBP) fused at the N-terminus of the C450CCMC sequence and a His_6_-tag attached to the C-terminus. The protein was purified to homogeneity (Figure 1F) by affinity chromatography on amylose agarose and on Ni-NTA resin, followed by anion exchange chromatography and gel filtration after cleaving the MBP tag. Although the final yields of purified protein were low (a few hundred micrograms from 6 L of bacterial culture), they were sufficient for functional characterization of the protein in vitro.

### The activity of Munc13-1 C450CCMC is autoinhibited by the N-terminal region and is stimulated by RIM and calmodulin

We next used well-established fusion assays with reconstituted proteoliposomes to compare the ability of the Munc13-1 C450CCMC fragment to support liposome fusion with that of the CCMC fragment corresponding to the conserved C-terminal region of Munc13-1, which has been extensively studied with these assays (e.g. refs. ^22,23,39^). These assays monitor both lipid mixing and context mixing between liposomes containing reconstituted synaptobrevin (V-liposomes) and liposomes containing syntaxin-1 and SNAP-25 (T-liposomes) using a membrane-bound fluorescence resonance energy transfer (FRET) pair and a soluble FRET pair, respectively. The T-liposomes are first incubated with Munc18-1, NSF and αSNAP to disassemble syntaxin-1-SNAP-25 complexes and form syntaxin-1-Munc18-1 complexes, and then they are incubated with the V-liposomes and the desired Munc13-1 fragment to induce liposome fusion. Liposome fusion is strongly stimulated by Ca^2+^ because it activates the Munc13-1 C_2_B domain, but some Ca^2+^-independent fusion may also be observed depending on the activity of the liposomes and added proteins.^22,53^

As expected from previous studies,^22^ there was no liposome fusion in the absence of any Munc13-1 fragment (Figure 2A,B, green curves), whereas the CCMC fragment spanning the Munc13-1 C-terminal region supported highly efficient, Ca^2+^-dependent liposome fusion (Figure 2A,B, black curves; see quantification at 400 s in Figure 2C,D). In contrast, the C450CCMC Munc13-1 fragment that in addition to the C-terminal region includes the C_2_A and CaMb domains was much less active (Figure 2A,B, red curves). The difference in the activity of the two fragments was particularly overt when comparing the amount of content mixing observed at 400 s, just 100 s after Ca^2+^ addition (Figure 2D), as the initial slope of the reaction was very steep for CCMC while there was a lag in the beginning of the reaction for C450CCMC (Figure 2B).

**Figure 2.**
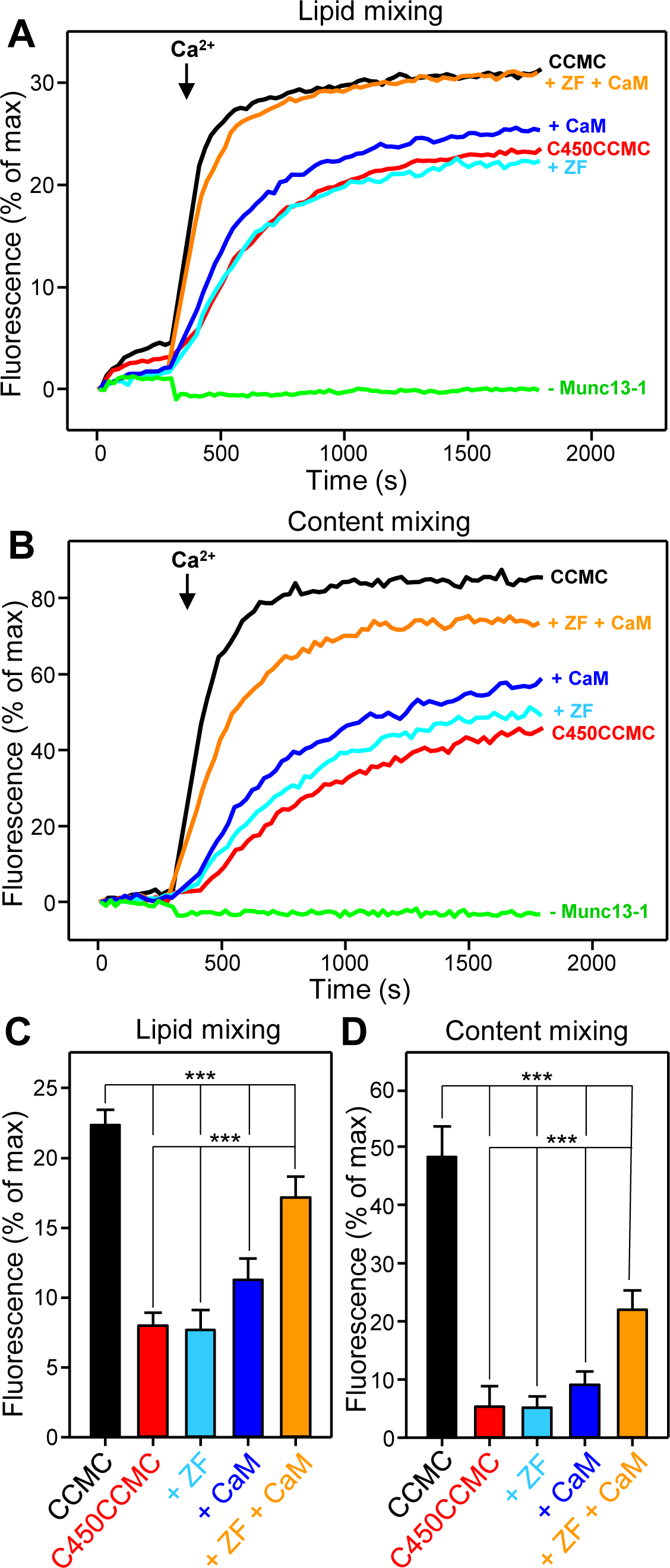
The Munc13-1 C450CCMC fragment is much less active than the Munc13-1 C-terminal region in liposome fusion assays and its activity is stimulated by the RIM2α ZF domain and calmodulin. (A, B) Lipid mixing between V- and T-liposomes (A) was measured from the fluorescence de-quenching of Marina Blue-labeled lipids and content mixing (B) was monitored from the development of FRET between PhycoE-Biotin trapped in the T-liposomes and Cy5-Streptavidin trapped in the V-liposomes. The assays were performed in the presence of Munc18-1, NSF, αSNAP and the Munc13-1 C-terminal region (CCMC, black curves) or the Munc13-1 C450CCMC fragment alone (red curves) or together with the RIM2α ZF domain (cyan curves) or calmodulin (CaM) (dark blue curves) or both (orange curves). Green curves correspond to a control experiments performed in the absence of any Munc13-1 fragment. Experiments were started in the presence of 100 μM EGTA and 5 mM streptavidin, and Ca^2+^ (600 μM) was added after 300 s. (C, D) Quantification of lipid mixing (C) and content mixing assays (D) performed in triplicate under the conditions of panels (A, B). Bars represent averages of the normalized fluorescence intensities observed at 400 s (i.e. 100 s after Ca^2+^ addition). Error bars represent standard deviations. Statistical significance and p values were determined by one-way analysis of variance (ANOVA) with the Holm-Sidak test (*** p < 0.001).

It is important to note that, in the crystal structure of the CCM Munc13-1 fragment, the N-terminal helix is oriented toward the MUN domain and that residues Asn1128 and Phe1131, which are important to open syntaxin-1,^25^ are located in the middle of the MUN domain. Hence, the sequences preceding the N-terminal helix in C450CCMC might fold back onto the MUN domain, potentially inhibiting its activity in catalyzing SNARE complex assembly. These observations and the low activity of the C450CCMC fragment in the liposome fusion assays led us to hypothesize that autoinhibitory interactions involving the CaMb domains and/or perhaps the C_2_A domain hinder the function of the MUN domain. Furthermore, we hypothesized that these autoinhibitory interactions might be released by calmodulin and the αRIM ZF domain, respectively. We tested these hypotheses by performing liposome fusion assays in the presence of C450CCMC plus calmodulin, the RIM2α ZF domain or both. Individually, the RIM2α ZF domain had no marked effect on liposome fusion, whereas calmodulin appeared to increase the activity of C450CCMC to a small extent that did not reach statistical significance (Figure 2A,B, light blue and dark blue curves, respectively; Figure 2C,D). However, liposome fusion was strongly enhanced when both calmodulin and the RIM2α ZF domain were added together with C450CCMC. These results support the notion that both the C_2_A domain and the CaMb domain inhibit Munc13-1 activity, and suggest that there is a synergy between RIM and calmodulin in relieving the inhibition caused by the N-terminal domains.

### The Munc13-1 CaMb domain binds to the MUN domain and calmodulin releases the interaction

The fact that the CaMb domain is adjacent to the Munc13-1 C-terminal region and is expected to be oriented toward the central sequences of the MUN domain that are key for syntaxin-1 opening (Figure 1A,B) suggests that the activity of the C-terminal region in the liposome fusion assays caused by the CaMb domain may arise from a direct intramolecular interaction between the two domains. To test this possibility, we turned to NMR spectroscopy, taking advantage of the power of two-dimensional (2D) ^1^H-^15^N heteronuclear single quantum coherence (HSQC) spectra to analyze protein-protein interactions.^54^ These spectra contain one cross-peak for the amide group of each non-proline residue in a uniformly ^15^N-labeled protein, and perturbations (cross-peak shifts or broadening) caused by addition of an unlabeled protein provide direct evidence for interaction between the two proteins. Thus, we used this approach to test whether the Munc13-1 MUN domain binds to a uniformly ^15^N-labeled Munc13-1 fragment that includes the CaMb domain (residues 450-492). Preparation of this fragment in isolation was hindered by its tendency to aggregate,^52^ but we were able to obtain this fragment in soluble form with a His_6_-tag by co-expressing it with calmodulin, binding the resulting complex to an Ni-NTA resin, removing calmodulin in the presence of 8 M urea and cleaving the His_6_-tag.

The ^1^H-^15^N HSQC spectrum of the Munc13-1 CaMb domain exhibited poor dispersion in the ^1^H dimension (Figure 3A), which is common in unstructured polypeptides. In addition to cross-peaks corresponding to Asn and Gln side chains in the upper right corner of the spectrum and to cross-peaks from Trp side chains in the lower left corner, about 35 backbone cross-peaks could be distinguished in the spectrum out 42 expected. The absence of some cross-peaks and the low intensities of some of the observed cross-peaks suggest that part of the sequence of the CaMb domain is aggregating or undergoing conformational exchange that broadens the cross-peaks, but a major part of the domain exhibits sharp cross-peaks characteristic of flexible sequences. Addition of the MUN domain caused small shifts in some of the ^1^H-^15^N HSQC cross-peaks of the CaMb domain and broadening beyond detection of other cross-peaks (Figure 3B). These results show that the Munc13-1 MUN domain indeed binds to the CaMb domain. The interaction between the two separate domains might be of moderate affinity, as these experiments were performed with 20 μM CaMb domain and 25 μM MUN domain, but intramolecular binding is expected to be considerably strengthened by localization of the CaMb domain near the MUN domain in native Munc13-1.

**Figure 3.**
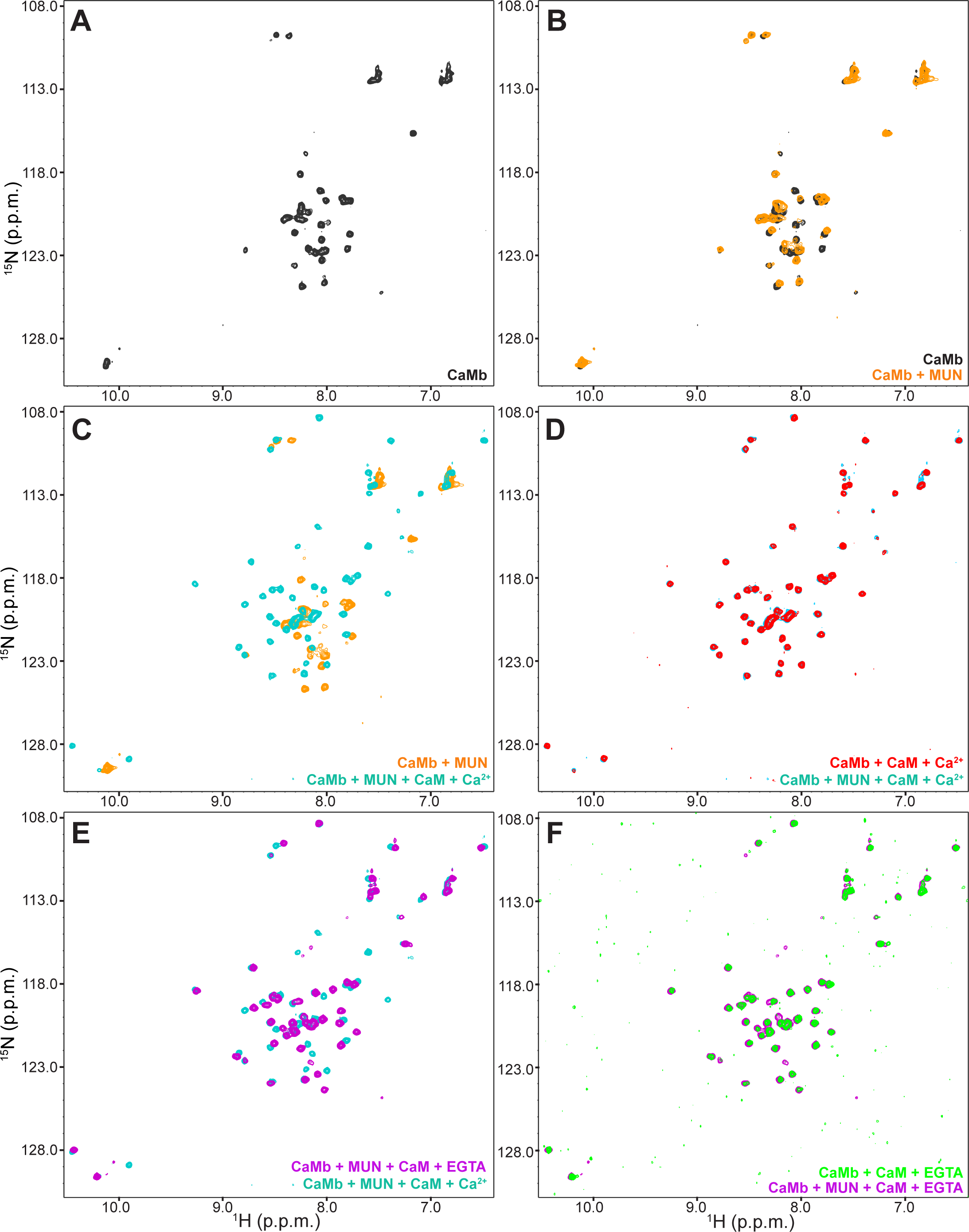
The Munc13-1 CaMb domain binds to the MUN domain and calmodulin competes with this interaction. (A-F) ^1^H-^15^N HSQC spectra of 20 μM ^15^N-labeled Munc13-1 CaMb domain: alone (A); alone (black contours) or in the presence of 25 μM MUN domain (orange contours) (B); in the presence of 25 μM MUN domain (orange contours) or 25 μM MUN domain plus 25 μM calmodulin (CaM) and 5 mM Ca^2+^ (blue contours) (C); in the presence of 25 μM calmodulin and 5 mM Ca^2+^ with (blue contours) or without (red contours) 25 μM MUN domain (D); in the presence of 25 μM MUN domain plus 25 μM calmodulin and 5 mM Ca^2+^ (blue contours) or 7.5 mM EGTA (purple contours) (E); and in the presence of 25 μM calmodulin and 7.5 mM EGTA with (purple contours) or without (green contours) 25 μM MUN domain (F).

We next tested whether Ca^2+^-bound calmodulin can compete with the MUN domain for binding to the CaMb domain by acquiring ^1^H-^15^N HSQC spectra of ^15^N-lalebed CaMb domain in the presence of unlabeled Ca^2+^-calmodulin and MUN domain. The resulting spectrum was dramatically different from that observed in the presence of MUN domain only and resembled the ^1^H-^15^N HSQC spectrum of a shorter ^15^N-CaMb fragment bound to calmodulin described earlier,^52^ with the excellent dispersion expected for a well-structured CaMb domain due to complex formation with calmodulin (Figure 3C). The much higher quality of this spectrum, compared to that observed for the CaMb domain bound to the MUN domain, and its similarity to the ^1^H-^15^N HSQC spectrum of the CaMb domain bound to Ca^2+^-calmodulin in the absence of MUN domain (Figure 3D), show that the CaMb domain/Ca^2+^-calmodulin complex does not bind to the MUN domain, which would have caused strong cross-peak broadening and perhaps shifts in observable cross-peaks.

We also acquired ^1^H-^15^N HSQC spectra of the CaMb domain in the presence of calmodulin, MUN domain and EGTA. The spectra still exhibited sharp, well-dispersed cross-peaks reflecting binding to calmodulin and multiple cross-peaks were in the same positions as in the spectra acquired in the presence of Ca^2+^, but some of the well-dispersed cross-peaks observed in the presence of Ca^2+^ disappeared and new cross-peaks appeared closer to the center of the spectrum (Figure 3E). These results are reminiscent of the previous observation that titration of the complex between the shorter CaMb fragment and Ca^2+^-calmodulin with EGTA led to an intermediate complex in which the CaMb fragment was released from the N-terminal domain of calmodulin but not from the C-terminal domain, which has a higher Ca^2+^ affinity.^52^ We note that we did not add Ca^2+^ or EGTA during the purification of calmodulin, but the C-terminal domain may have captured Ca^2+^ from the media during purification. Therefore, the ^1^H-^15^N HSQC spectrum of the CaMb domain in the presence of MUN, calmodulin and EGTA likely corresponds to the intermediate complex with the CaMb domain bound only to the calmodulin C-terminal domain. This spectrum was again very similar to that of the CaMb domain in the presence of calmodulin and EGTA but in the absence of MUN domain (Figure 3F), showing that the intermediate CaMb/calmodulin complex does not bind to the MUN domain.

To analyze the interplay between calmodulin, the CaMb domain and the MUN domain in the context of a Munc13-1 fragment containing both domains such that their interaction would most likely be intramolecular, we performed gel filtration experiments with a fragment spanning the CaMb, C_1_, C_2_B and MUN domains (residues 450-1531; referred to as 450CCM) (Figure 4). Comparison of the elution profiles observed for calmodulin, 450CCM and an equimolar mixture of them in the presence of 5 mM EGTA in the elution buffer, and analysis of the fractions by SDS-PAGE, showed that most of the calmodulin of the mixture eluted at a similar volume as isolated calmodulin, albeit a little bit earlier, and that a small amount of calmodulin co-eluted with 450CCM (Figure 4A). In contrast, most of the calmodulin co-eluted with 450CCM in analogous experiments performed in the presence of Ca^2+^ (Figure 4B). These results support the notion that, as reported previously^52^, calmodulin binds tightly to the Munc13-1 CaMb domain in the presence of Ca^2+^ but retains binding of weak affinity in the absence of free Ca^2+^ or at very low Ca^2+^ concentrations such as those present in the cytoplasm of presynaptic terminals at rest. Moreover, our NMR data show that binding of calmodulin to the Munc13-1 CaMb domain competes with the interaction between the CaMb and MUN domains, supporting the hypothesis that such an interaction inhibits Munc13-1 activity and that calmodulin releases this inhibition by binding to the CaMb domain.

**Figure 4.**
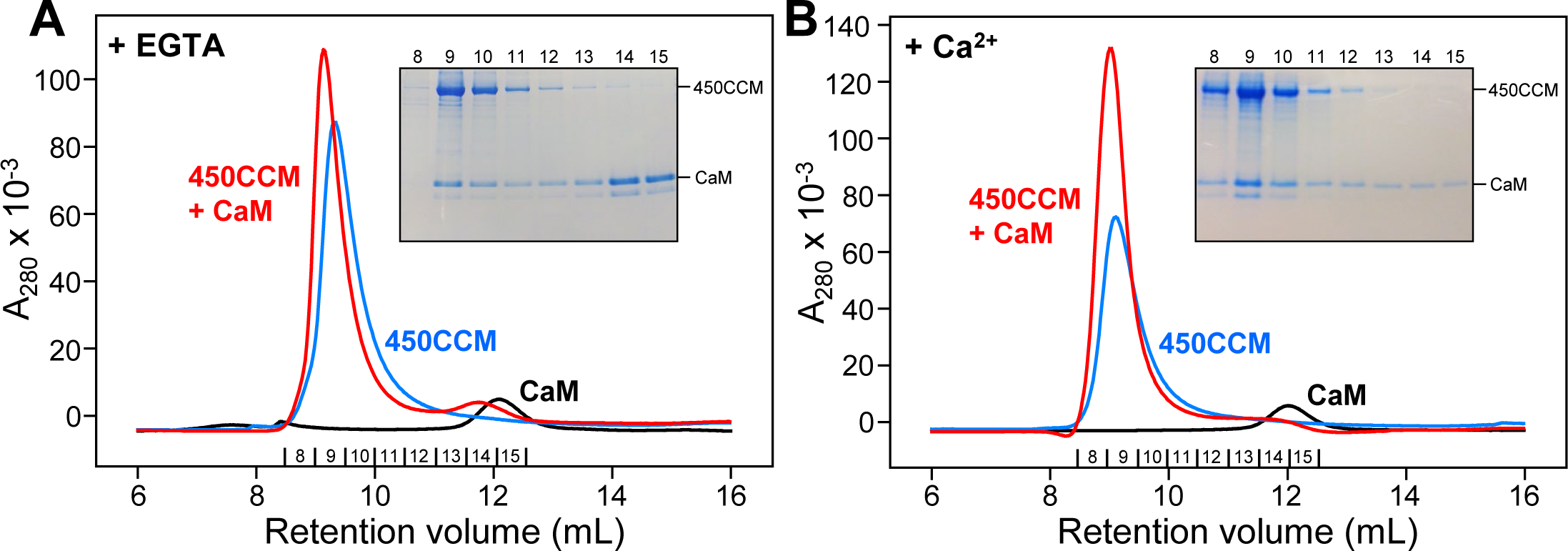
The Munc13-1 450CCM fragment binds to calmodulin more tightly in the presence of Ca^2+^ than in the presence of EGTA. (A,B) Size exclusion chromatography (Superdex^TM^ 75 300/10 GL) profiles of Munc13-1 450CCM (blue curves), His_6_-calmodulin (CaM) (red curves) and 450CCM plus His_6_-calmodulin (red curves) in the presence of 5 mM EGTA (A) or 5 mM Ca^2+^ (B). The insets show SDS-PAGE analyses with Comassie blue staining of the fractions indicated below the chromatograms.

### The Munc13-1 MUN domain binds to the C_2_A domain homodimer but not to the C_2_A domain/RIM ZF heterodimer

The inhibition of the activity of the Munc13-1 C-terminal region by the C_2_A domain observed in our liposome fusion assays might in principle arise because homodimerization of the C_2_A domain may lead to opposite relative orientations of the two C-terminal regions of the homodimer, which may thus hinder each other. However, it is also plausible that the inhibition arises because of direct interactions of the C_2_A domain with the MUN domain that hinder its activity. To test the second possibility, we also used NMR spectroscopy. The Munc13-1 C_2_A domain homodimer (34 kDa) yields poor quality 2D ^1^H-^15^N HSQC spectra^44^ and the quality is even poorer for the considerably larger MUN domain (73 kDa; unpublished results). Since the first trace of ^1^H-^15^N HSQC spectra (referred to as 1D ^1^H-^15^N HSQC spectra) yields good signal sensitivity even when the quality of the 2D spectrum is poor, and the intensity of the overall resonance envelope is expected to decrease upon binding to an unlabeled protein because of resonance broadening (e.g. ref. ^24^), we used this approach to monitor binding of the Munc13-1 MUN domain to ^15^N-labeled C_2_A domain.

Indeed, we observed a strong decrease in the intensity of the signal envelope in 1D ^1^H-^15^N HSQC spectra of the C_2_A domain upon addition of MUN domain (Figure 5A,B). However, when we added the RIM2α ZF domain in addition to the MUN domain, the envelope signal intensity was actually increased with respect to that of the C_2_A domain alone (Figure 5C) and the spectrum was comparable to that observed for the C_2_A domain bound to the RIM2α ZF domain in the absence of the MUN domain (Figure 5D). This increased signal intensity arises because of the smaller size of the C_2_A domain/ZF heterodimer compared to that C_2_A domain homodimer. These results show that the Munc13-1 C_2_A domain binds to the MUN domain and that formation of the C_2_A domain/ZF heterodimer releases this interaction, supporting the hypothesis that interactions of the C_2_A domain with the MUN domain inhibit Munc13-1 activity and that this inhibition is relieved by RIM.

**Figure 5.**
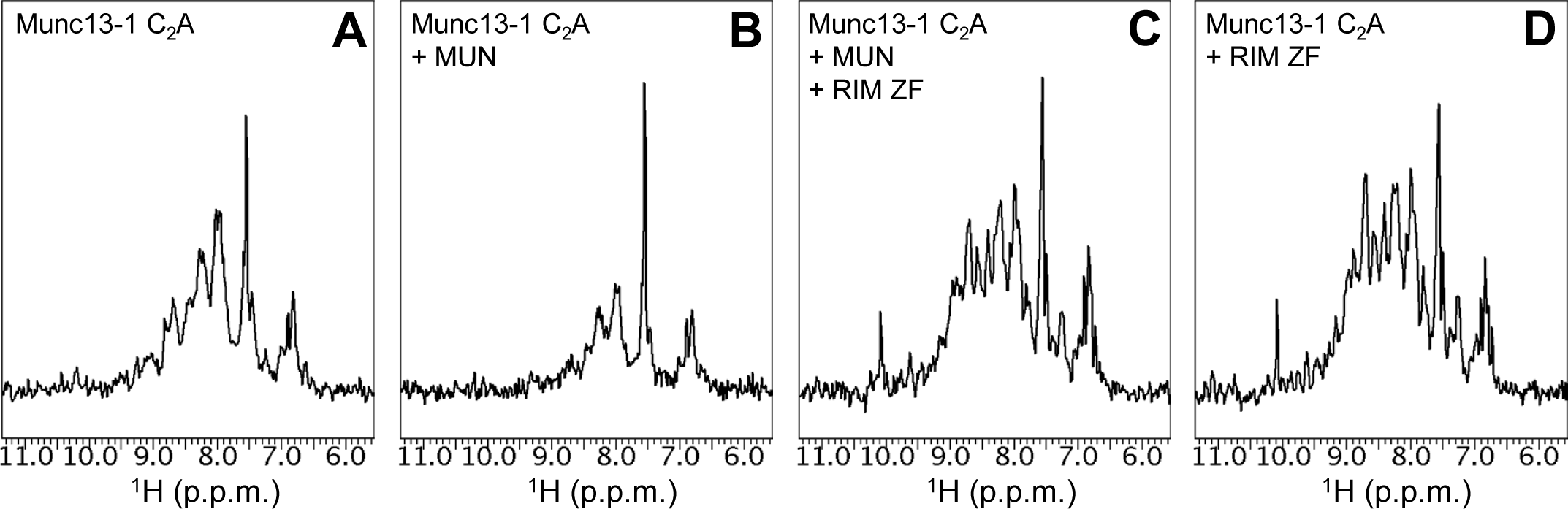
The Munc13-1 C_2_A domain binds to the MUN domain and the RIM2α ZF domain competes with this interaction. (A-D) 1D ^1^H-^15^N HSQC spectra of 50 μM ^15^N-labeled Munc13-1 C_2_A domain alone (A) or in the presence of 50 μM MUN domain (B), 50 μM MUN domain and 50 μM RIM2α ZF domain (C), or 50 μM RIM2α ZF domain (D).

## Discussion

The basic steps that mediate synaptic vesicle priming and neurotransmitter release are reasonably well understood, but the molecular mechanisms underlying the regulation of neurotransmitter release during a wide variety of presynaptic plasticity processes that are crucial for information processing in the brain are largely unknown. Because Munc13-1 plays a central role in promoting SNARE complex assembly and in addition acts as a master regulator of release during diverse forms of presynaptic plasticity, it is particularly important to elucidate the mechanisms of action of the multiple domains of this large active zone protein and how they are integrated. The conserved C-terminal region of Munc13-1, which constitutes the ‘executive’ part of the molecule, has been extensively studied and shown to play a key role in SNARE complex assembly by helping to open syntaxin-1^24,25^ while bridging the vesicle and plasma membranes.^22,23^ Moreover, the notions that the Munc13-1 C-terminal region can bridge the two membranes in two different orientations^39^ that underlie two primed states with distinct release probabilities^42,43^, and that DAG and Ca^2+^ favor the more active orientation^41^, have provided an attractive model for Ca^2+^- and DAG-dependent presynaptic plasticity mediated by this region. However, the molecular mechanisms underlying the functions of the variable N-terminal region of Munc13-1 and presynaptic plasticity processes involving this region are much less well understood, in part because of the difficulty of expressing and purifying full-length Munc13-1 and large fragments including N-terminal domains. We have now overcome these problems and purified the Munc13-1 C450CCMC fragment, which contains the C_2_A and CaMb domains in addition to the C-terminal region. We show that this fragment exhibits a markedly lower activity than the CCMC fragment spanning the C-terminal region in liposome fusion assays, and that the activity of C450CCMC is stimulated by calmodulin and the RIM2α ZF domain in what appears to be a synergistic manner. Moreover, we provide evidence supporting the notion that autoinhibitory interactions of the C_2_A and CaMb domains with the MUN domain control Munc13-1 activity and that these inhibitory interactions are relieved by RIM and calmodulin, respectively (Figure 6). Overall, these results suggest that an intricate set of intramolecular and intermolecular interactions modulate the function of Munc13-1 in neurotransmitter release and govern varied forms of presynaptic plasticity.

**Figure 6.**
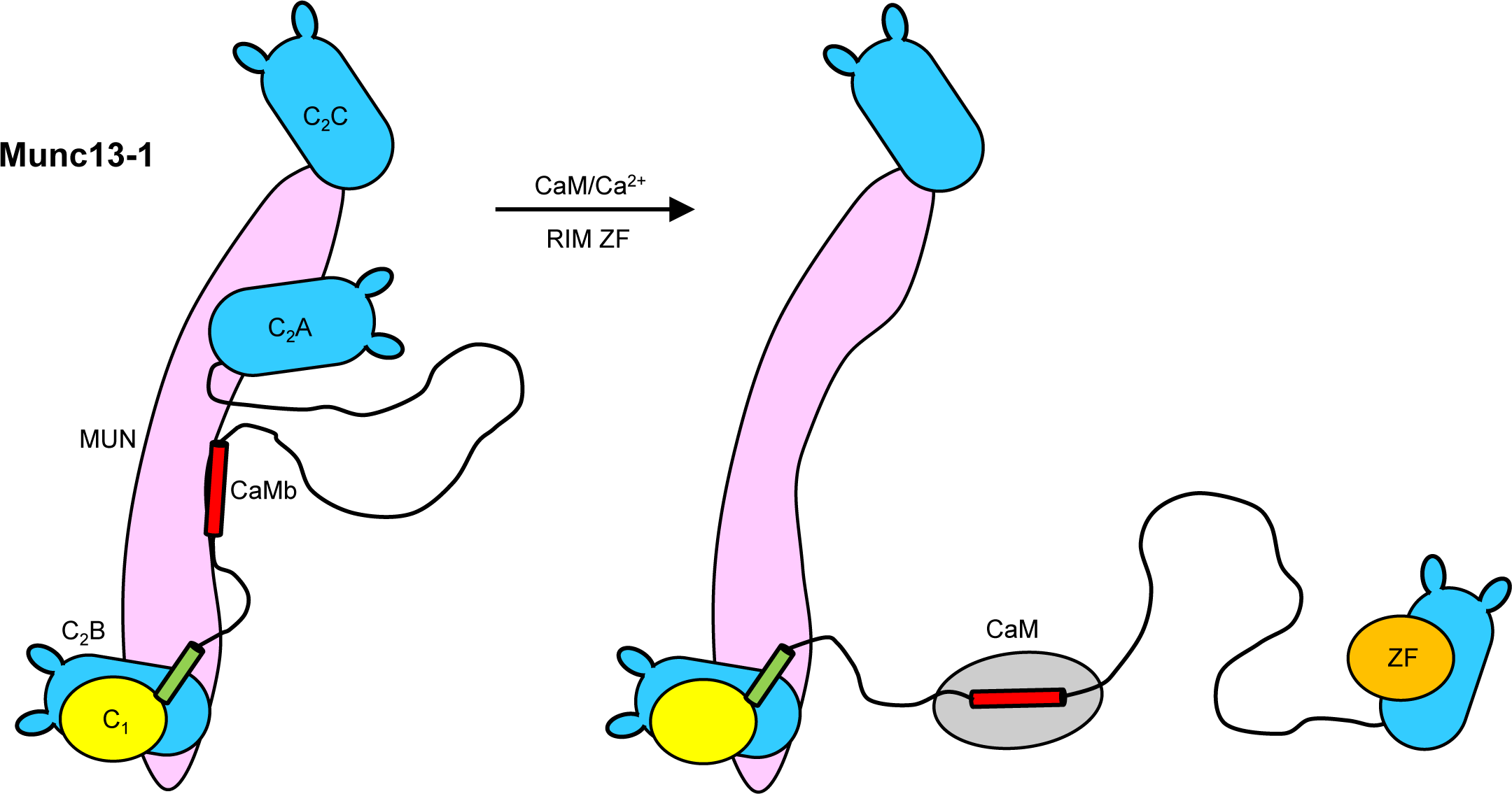
Model of how the Muc13-1 N-terminal region, calmodulin and RIM control Munc13-1 activity. The model predicts that the Munc13-1 C_2_A and CaMb domains bind intramolecularly to the MUN domain, inhibiting its activity in promoting SNARE complex assembly and synaptic vesicle priming. Binding of the RIM ZF domain (ZF) to the C_2_A domain and of calmodulin (CaM) to the CaMb domain relieves the inhibitory interactions. Calmodulin is postulated to perform this action more efficiently when Ca^2+^ concentrations increase in the presynaptic terminal, but may also activate Munc13-1 to a lower extent at basal Ca^2+^ levels. Before activation by RIM, the Munc13-1 C_2_A domain forms a homodimer (not illustrated in the figure), which depending on its geometry may hinder the membrane-membrane bridging activity of the Munc13-1 C-terminal region. Inhibition of this activity could also involve interactions of the C_2_A domain with the C_1_/C_2_B region or the C_2_C domain, but this possibility would require a different arrangement of the C_2_A domain from that shown on the left diagram.

The observation that the presence of the C_2_A and CaMb domains in C450CCMC inhibits the activity of the Munc13-1 C-terminal region in liposome fusion assays (Figure 2) correlates with previous observations showing that these two N-terminal domains can play inhibitory functions that are relieved by αRIMs or calmodulin, respectively. Thus, on one hand, functional studies showed that binding of Ca^2+^-calmodulin to Munc13-1 or the ubiquitous Munc13-2 isoform ubMunc13-2 enhances priming of synaptic vesicles and of chromaffin granules,^49,55^ indicating that Ca^2+^-calmodulin somehow activates Munc13s containing a CaMb domain. Moreover, the observation that deletion of residues 1-520 of Munc13-1 that span the entire N-terminal region causes no overt effect on Ca^2+^-evoked neurotransmitter release and a moderate impairment of vesicle priming, whereas both evoked release and priming are strongly impaired by deletion of only the C_2_A domain,^51^ suggested that some sequence within residues 151-520 inhibits Munc13-1 activity and the C_2_A domain plays a role in relieving this inhibition. On the other hand, Munc13-1 C_2_A domain homodimerization also plays an inhibitory function, as the strong priming defect observed in the absence of αRIMs can be rescued by an N-terminal fragment of RIM1α containing the ZF domain that breaks the Munc13-1 C_2_A domain homodimer to form a heterodimer, and this priming defect can also be rescued by ubMunc13-2 bearing a K32E mutation that precludes homodimerization but not by wild type ubMunc13-2.^50^ These observations might suggest that the only function of the αRIM-Munc13-1 interaction is to relieve the inhibition caused by Munc13-1 C_2_A domain homodimerization. However, a double point mutation in the C_2_A domain (E128K,E137K) that abrogates αRIM binding strongly impairs vesicle priming and evoked release, and these defects are partially alleviated by including the K32E mutation that precludes homodimerization to yield a K32E,E128K,E137K triple mutant, but substantial defects in priming and evoked release remain.^51^ These findings show that αRIM-Munc13-1 binding has an important function beyond disruption of the C_2_A domain homodimerization, which is likely associated to a role in helping to localize Munc13-1 at the active zone.^51^

This role and other aspects of the functions played by the Munc13-1 N-terminal region cannot be reproduced with the limited number of components used in our reconstitution assays, but the observation that inclusion of the RIM2α ZF domain and calmodulin strongly stimulates the activity of C450CCMC in the liposome fusion assays (Figure 2) shows that fundamental features of the function of the Munc13-1 N-terminal region are indeed recapitulated in these assays. These results add to multiple correlations between liposome fusion and physiological data that we have established over the years with this reconstitution system^4,6,16,22,23,39,41,53^ and further illustrate the usefulness of this system to study the mechanisms of action of the neurotransmitter release machinery. We do note that the activity of C450CCMC was not as high as that of CCMC even upon activation by the RIM2α ZF domain and calmodulin (Figure 2). However, this is not surprising because in neurons RIMs form part of the active zone at the presynaptic plasma membrane and, as described above, they help to localize Munc13s to the active, a role that cannot be played in our liposome fusion assays by the soluble RIM2α ZF domain.

Our NMR data revealing direct interactions of the Munc13-1 C_2_A and CaMb domains with the MUN domain that are relieved by the RIM2α ZF domain and calmodulin, respectively (Figure 3, 5), suggest that these interactions underlie the inhibitory functions of the C_2_A and CaMb domains, likely because they hinder the function of MUN domain in opening syntaxin-1. Moreover, the observation that both the RIM2α ZF domain and calmodulin are required to strongly enhance the activity of C450CCMC in the liposome fusion assays suggests that there may be some synergy in the actions αRIMs and calmodulin. There is also likely to be an interplay between the interactions involving the N-terminal domains and other activities of the Munc13-1 C-terminal region, which plays a crucial role in neurotransmitter release not only by helping to open syntaxin-1 ^24,25^ but also by bridging the vesicle and plasma membranes.^22,23^ Thus, accumulation of Ca^2+^ during repetitive stimulation mediates short-term presynaptic plasticity in part through the Munc13-1 CaMb domain^49^, which probably controls the syntaxin-1 opening function, and in part by activating the C_2_B domain^37^, which is involved in membrane-membrane bridging^22,23^. Based on the two-primed state model,^39,42^ Ca^2+^ binding to the C_2_B domain is expected to facilitate the slanted orientation of Munc13-1 with respect to the plasma membrane that is believed to be present in the primed state with the higher release probability.^41^ The C_2_B domain is activated with an apparent K_D_ for Ca^2+^ between 600 and 950 nM in liposome fusion assays^41^, whereas the action of calmodulin on the CaMb domain may involve two components operating also at submicromolar, albeit distinct, Ca^2+^ concentrations through its two domains.^52^ Our NMR experiments suggest that calmodulin can displace the CaMb domain from the MUN domain not only when calmodulin is saturated with Ca^2+^ but also when calmodulin binds to the CaMb domain through only its C-terminal domain (Figure 3), which has very high Ca^2+^ affinity^52^ and may bind to Munc13-1 even at basal Ca^2+^ concentrations. However, the effective affinity of the CaMb domain for the MUN domain is likely to be dramatically enhanced in the context of full-length Munc13-1 due to their expected proximity (Figure 1B); therefore, calmodulin may not be able to compete with this intramolecular interaction at basal Ca^2+^ concentrations or at least not as effectively as when Ca^2+^ binds also to the calmodulin N-terminal domain, as suggested by our gel filtration data (Figure 4).

Further experiments, including structural studies of C450CCMC or similar Munc13-1 fragments, will be required to test these ideas. However, our results and the already available structure of the CCM fragment (Figure 1B) suggest that an intricate set of intramolecular interactions and intermolecular interactions with binding partners coordinate the functions of the various Munc13-1 domains. This would not be surprising, as complex sets of interactions also govern other aspects of the molecular mechanisms that underlie neurotransmitter release, including the orchestration of SNARE complex assembly by Munc18-1 together with Munc13-1^4,15,21^, or the interplay between synaptotagmin-1, complexin and the SNAREs in mediating fast Ca^2+^-induced synaptic vesicle fusion^27,29–31^. There is little doubt that the neurotransmitter release machinery functions as a well-oiled engine with clockwork precision.

## Materials and Methods

### Protein expression and purification

Expression and purification of full-length Homo sapiens SNAP-25A (with its four cysteines mutated to serine), full-length Rattus norvegicus synaptobrevin-2, full-length Rattus norvegicus Munc18-1, full-length Cricetulus griseus NSF V155M mutant, full-length Bos taurus α-SNAP, full-length Rattus norvegicus syntaxin-1A and rat RIM2α ZF domain (residues 82-142) in E. coli were described previously^4,17,46,56,57^.

A pProEx2 vector to express human calmodulin (CALM2) with a His_6_ tag and a TEV cleavage site at the N-terminus^58^ was a kind gift from Vincent S. Tagliabracci. BL21(DE3) cells containing pProEx2-CALM2 were grown until OD_600_ ∼0.6 in the presence of 100 μg/ml ampicillin and induced overnight at 22 °C with 0.4 mM isopropyl-B-D-thiogalactopyranoside (IPTG). Cells were harvested by centrifugation and resuspended in ice-cold lysis buffer 50 mM Tris pH 8.0, 250 mM NaCl, 20 mM Imidazole, protease Inhibitor cocktail from Sigma-Aldrich and 0.1% β-mercaptoethanol (ME). Cells were broken by passing four times through an Avestin (Ottawa, Canada) cell disrupter at > 12,000 p.s.i. After centrifugation of the lysate for 40 min at 39,191 g at 4°C, the supernatant was bound to a Ni-NTA resin (1ml bead column/L culture) and washed with 10 column volumes (C.V.) of 50 mM Tris pH 8.0, 1M NaCl, 20 mM Imidazole 0.1% ME followed by 10 C.V. of lysis buffer. The protein was eluted with 50 mM Tris pH 8.0, 250 mM NaCl, 1 mM TECEP, 300 mM Imidazole, and dialized overnight in the same buffer without Imidazole. The protein was concentrated and further purified on gel filtration (Superdex^TM^ 75 300/10 GL) equilibrated with 50 mM Tris pH 8.0, 250 mM NaCl, 1 mM TCEP. For NMR experiments, the His_6_-tag was removed by cleavage with His_6_-TEV and the protease was removed with Ni-NTA resin.

All plasmids used to express recombinant Munc13-1 fragments originated from a vector encoding full-length rat Munc13-1 (NM-022861) used to rescue Munc13 function (e.g. ref. ^22^) in neurons that was kindly provided by Christian Rosenmund, and were generated by standard molecular biology techniques with custom designed primers. All Munc13-1 fragments that included the MUN domain contained a deletion in a large variable loop (residues 1408-1452) that improves the solubility.^24^ Expression and purification of Munc13-1 MUN domain (residues 859-1531, Δ1408-1452) and CCMC (residues 529-1735, Δ1408-1452) in bacteria was described earlier.^23,38^ A vector to express the Munc13-1 450CCM fragment (residues 450-1531, Δ1406-1463) was constructed into pFASTBacHTb vector. The construct was used to generate a baculovirus using the Bac-to-Bac system (Invitrogen; Waltham, MA). Sf9 insect cells were infected with the baculovirus, harvested about 72–96 hr post-infection, and re-suspended in lysis buffer (50 mM Tris pH8.0, 250 mM NaCl, 10 mM imidazole, 1mM TCEP). Cells were lysed and centrifuged at 30,966 rpm for 45 min. The clear supernatant was incubated with Ni-NTA resin at 4°C for 2 hr, then the beads were washed with: (i) lysis buffer; (ii) lysis buffer containing 1% Triton X-100; (iii) lysis buffer containing 1M NaCl; and (iv) lysis buffer. The protein was eluted in lysis buffer with 50-400 mM imidazole, incubated with TEV to remove the His-tag, and purified by ion exchange chromatography (Hitrap^TM^ Q HP Cytiva).

A plasmid to express the Munc13-1 CaMb domain (residues 450-492) was constructed into a pETDuet vector. A His_6_-tag followed by a TEV cleavage site was linked to the N-terminus of Munc13-1(450-492). Calmodulin was cloned into a second cloning site of the pETDuet vector. His_6_-Munc13-1(450-492) and calmodulin were co-expressed in E.Coli BL21 (DE3) cells grown to an OD_600_ of 0.8-1.0 and induced overnight at 20 °C with 0.4mM IPTG. Before induction, 2 mM CaCl_2_ were added to the cells. Cells were harvested by centrifugation and re-suspended in re-suspension buffer (50 mM HEPES pH 7.4, 300 mM NaCl) prior to lysis. Cell lysates were centrifuged at 48,000 x g for 40 min. Clarified supernatant was flowed through Ni-NTA resin twice (1 ml of beads/1 L of cell culture). The Ni-NTA resin was incubated with denaturation buffer (50 mM HEPES pH7.4, 300 mM NaCl, 8 M Urea, 2 mM EGTA, 2 mM MgCl_2_) at RT for 30 min and was washed with 3 C.V. of denaturation buffer to remove calmodulin, followed by washing with 6 C.V. washing buffer (50 mM HEPES pH7.4, 300 mM NaCl, 2 mM EGTA, 2 mM MgCl_2_). His_6_-Munc13-1(450-492) was eluted from Ni-NTA resin with 4 C.V. elution buffer (50 mM HEPES pH 7.4, 300 mM NaCl, 2 mM EGTA, 2 mM MgCl_2_, 0.5 M Imidazole) and further purified with size exclusive chromatography (Superdex^TM^ 75 300/10 GL) in gel filtration buffer (25 mM HEPES pH7.4, 125 mM NaCl). The His_6_ tag was removed with His6-TEV cleavage and the protease was removed by flowing through Ni-NTA resin, yielding the purified Munc13-1(450-492) fragment.

A plasmid to express the Munc13-1 C450CCM fragment (residues 1-1735, Δ151-449, Δ1408-1452) preceded by MBP and a TEV cleavage site at the N-terminus and followed by a His_6_-tag at the C-terminus was generated in a pMal-C2 vector. The protein was expressed in E.Coli BL21 (DE3) cells in LB medium. When the OD_600_ reached 0.6-0.8, expression was induced with 0.3 mM IPTG overnight at 18 °C. The cells were centrifuged 4,000 rpm for 30 min, re-suspended in re-suspension buffer (50 mM Tris pH 8, 250 mM NaCl, 20 mM Imidazol) containing Rocher cOmplete™ Protease Inhibitor Cocktail, 1mM TCEP and 5 mM EDTA, broken by passing four times through an Avestin cell disrupter at 12,000 p.s.i., and spun at 48,000 x g for 45 min. The supernatant was incubated with amylose agarose column and the resin was rotated at 4 °C for 10 min. The resin was extensively washed with re-suspendion buffer and eluted with 10 mM maltose in re-suspension buffer containing protease inhibitors and 1 mM TCEP, and loaded onto Ni-NTA resin. The resin was rotated at 4°C for 10 min, and washed with 10 C.V. each of re-suspension buffer, re-suspension buffer containing 1 M NaCl and re-suspension buffer. The protein was eluted with 6 C.V. of 200 mM imidazole. The MBP moiety was removed by cleavage with TEV protease (OD_280_ 1 of TEV for OD_280_ 20 of fusion protein). The protein was dialyzed overnight at 4 °C and purified by gel filtration on Superdex200 in 20 mM Tris pH 8, 250 mM NaCl, 1 mM TCEP.

### Liposome fusion assays

Lipid and content mixing assays were performed as previously described with slight modifications.^59^ Briefly, T-liposomes contained 38% 1-palmitoyl-2-oleoyl-sn-glycero-3-phosphocholine (POPC), 18% 1,2-dioleoyl-sn-glycero-3-phospho-L-serine (DOPS), 20% 1-palmitoyl-2-oleoyl-sn-glycero-3-phosphoethanolamine (POPE), 2% L-a-Phosphatidylinositol-4,5-bisphosphate (PIP_2_), 2% 1-palmitoyl-2-oleoyl-*sn*-glycerol (DAG), and 20% cholesterol. V-liposomes contained 40% POPC, 6.8% DOPS, 30.2% POPE, 20% Cholesterol, 1.5% 1,2-dipalmitoyl-*sn*-glycero-3-phosphoethanolamine-N-(7-nitro-2-1,3-benzoxadiazol-4-yl) (ammonium salt) (NBD-PE), and 1.5% 1,2-Dihexadecanoyl-*sn-*glycero-3-phosphoethanolamine (Marina Blue DHPE). T-Lipopsomes were prepared with syntaxin-1:SNAP-25:lipid ratio 1:5:1000, and V-liposomes with synatobrevin:lipid ratio 1:1000. In the final reactions, T-liposomes (250 mM total lipid) were first incubated with 1 μM Munc18-1, 0.2 μM NSF, 0.5 μM aSNAP, 2 mM ATP, 2.5 mM MgCl_2_, 5 mM streptavidin, and 100 μM EGTA for 25 min at 37°C, and then were mixed with V-liposomes (125 mM total lipid), 0.2 μM SNAP-25, and different Munc13-1 fragments at 0.05 μM concentration. Selected reactions also included 2 μM RIM2α ZF domain and/or 1 μM Calmodulin. After 5 min 0.6 mM Ca^2+^ was added to stimulate fusion, and 1% b-OG was added after 30 min to solubilize the liposomes. The fluorescence signals from Marina Blue (excitation at 370 nm, emission at 465 nm) and Cy5 (excitation at 565 nm, emission at 670 nm) were recorded to monitor lipid and content mixing, respectively. The lipid mixing data were normalized to the maximum fluorescence signal observed upon detergent addition. The content mixing data were normalized to the maximum Cy5 fluorescence observed after detergent addition in control experiments with only T and V liposomes without external streptavidin.

### NMR Spectroscopy

NMR spectra were acquired at 25 °C on an Agilent DD2 spectrometer operating at 600 MHz and equipped with a cold probe. Two-dimensional ^1^H-^15^N HSQC spectra were obtained on samples containing 20 μM ^15^N-Munc13-1 CaMb domain, with or without 25 μM Munc13-1 MUN domain and/or 25 μM Calmodulin in 25 mM HEPES pH 7.4, 125 mM NaCl, 0.5 mM TCEP, 8% D_2_O). For spectra containing Calmodulin, 5 mM CaCl_2_ or 7.5 mM EGTA were included in the NMR buffer. The spectra were processed with NMRPipe^60^ and visualized with NMRViewJ.^61^ One-dimensional ^1^H-^15^N HSQC spectra of 50 μM ^15^N-Munc13-1 C_2_A domain in the presence or absence of 50 μM MUN domain and/or 50 μM RIM2α ZF domain were acquired in 50 mM Tris pH 8.0, 150 mM NaCl, 10% D_2_O. The spectra were processed and visualized with Agilent vnmrJ software.

### Gel filtration binding assay

To analyze binding of calmodulin to Munc13-1 450CCM, 500 μl of 5 μM 450 CCMC alone, 5 μM His_6_-calmodulin alone, or 5 μM 450CCM incubated with 5 μM His_6_-calmodulin at RT for 15min were loaded on a SuperdexTM 75 300/10 GL column pre-equilibrated in 25mM HEPES pH7.4, 125 mM NaCl, 0.5 mM TCEP with 5 mM CaCl_2_ or 5mM EGTA. Fractions were analyzed by SDS-PAGE followed by Comassie blue staining.

## Acknowledgments

We thank Yibin Xu for technical help with protein preparation and NMR experiments with the Munc13-1 C_2_A domain. This work was supported by grant I-1304 from the Welch Foundation (to JR) and by NIH Research Project Award R35 NS097333 (to JR). J.R. holds the Virginia Lazenby O’Hara Chair in Biochemistry.

